# Bark beetles as ecosystem engineers: triggered tree mortality rearranges the assemblage of Tree-related Microhabitats in old-growth coniferous forest

**DOI:** 10.1101/2025.04.11.648382

**Authors:** Fabian Przepióra, Michał Ciach

## Abstract

Although species with ecosystem engineering capabilities like woodpeckers, ungulates or saproxylic insects may initiate the formation of Tree-related Microhabitats (TreMs), their overall impact on TreM assemblages remains unexplored. Bark beetle activity can increase tree mortality, accelerating the emergence of canopy gaps with altered tree species composition, stand structure and deadwood resources, all of which can influence TreM profiles and cascade on biodiversity. We investigated the TreM assemblage in a strictly-protected old-growth Norway spruce *Picea abies* forest in the Tatra Mountains (Carpathians, S Poland), comparing bark beetle-induced canopy gaps with undisturbed closed-canopy reference sites. Although TreM richness, density and diversity did not differ between the canopy gaps and reference sites, the frequency and density of woodpecker cavities, splintered stems, cracks, bark pockets, bark shelters, fruiting bodies of polypore fungi and epiphytic ferns was higher in the canopy gap TreM assemblage. In contrast, the frequency and density of exposed sapwood, shaded dead branches, dead tree tops, coarse bark, root buttress cavities, cankers, resin runs and microsoil in the tree crown or bark were higher in the reference sites. TreM richness, density and diversity were correlated with the density of living or dead standing trees and tree species richness, all related to previous bark beetle activity. Bark beetle-induced canopy gaps did not increase the total TreM abundance or diversity, but by hosting more saproxylic TreMs, locally altered the TreM composition. Our study underlines the importance of ecosystem engineering by bark beetles as drivers of the heterogeneity of temperate coniferous montane forests.

**Highlights:**

- TreMs in coniferous forests are abundant but less diverse than in deciduous ones
- Canopy gaps have a distinct TreM profile compared to intact old-growth forest
- Canopy gaps are hot-spots of certain TreMs in coniferous forests
- Saproxylic TreMs are 3–9 times more abundant in canopy gaps than in intact forests
- Tree mortality triggered by bark beetles increases habitat heterogeneity

## 1. Introduction

Tree-related Microhabitats (TreMs) provide numerous species with sites for development, foraging and shelter (Larrieu et al. 2018). Certain TreMs support a great variety of organisms: for example, tree cavities in the temperate forests of Europe are utilized by birds, bats, rodents, carnivores, amphibians, invertebrates and fungi (Nilsson and Baranowski 1997; Czeszczewik et al. 2008; Fritz and Heilmann-Clausen 2010; Piraccini et al. 2017; Van Der Hoek et al. 2017). Some TreMs may support fewer species, but these often include specialized and rare organisms, such as aquatic insects thriving in water-filled dendrotelmata or endangered flies dependent on sap runs (Petermann and Gossner 2022; Van Steenis 2023). While most TreMs are strongly associated with tree age, e.g. coarse bark on mature trees, some may be more commonly found on dead standing trees (hereafter, snag), e.g. fungal fruiting bodies, insect galleries or loose bark patches, or particular tree species, e.g. resin runs on spruces (Paillet et al. 2017; Kozák et al. 2023). TreMs also depend on rather rare events like hurricane-force winds or fire, which give rise to broken stems or charred wood, for example (Larrieu et al. 2022). TreM occurrence is also influenced by topography, e.g. the elevation or aspect of slopes (Przepióra et al. 2025), stand characteristics, e.g. tree size structure, tree species composition and the occurrence of injured trees or snags (Paillet et al. 2017; Larrieu et al. 2022), and the microclimate, which impacts on wood decomposition and the growth of epiphytic organisms (Remm and Lõhmus 2011; Man et al. 2022). All such factors mediating a TreM assemblage are often related with forest disturbances of varying severity and scale, including those induced by organisms (Zemlerová et al. 2023).

Ecosystem engineers significantly modify their environment, creating habitats and conditions that influence ecosystem structure, support other species and, therefore, increase biodiversity (Jones et al. 2010). Ecosystem engineers include species that influence TreM formation. For example, woodpeckers excavate breeding cavities and feeding holes (Larrieu et al. 2018), while deer or bears strip bark, thus exposing wood and producing sap runs that can result in fungal infections and trunk cavities (Zyśk-Gorczyńska et al. 2015; Broughton et al. 2022). The foraging activity of beavers promotes the growth of multi-trunk trees that bear TreMs such as mud and twig piles, or provide snags in which the exit holes of xylophagous insects are later found (Mourant et al. 2018; Przepióra and Ciach 2022). The role of invertebrates in TreM formation is amply illustrated by the great capricorn beetle *Cerambyx cerdo*, which excavates tunnels in wood that subsequently provide bats, amphibians and reptiles with shelter, hibernation or foraging sites (Gottfried et al. 2019; Borczyk et al. 2022). While some studies have focused on the link between specific TreMs and the animals that contribute to their formation, the overall impact on TreM assemblages of species, including the ones with ecosystem engineering capabilities, remains largely unexplored.

In temperate coniferous forests, bark beetles are recognized as key ecosystem engineers (Müller et al. 2008). By entering through the bark to lay eggs in the phloem, bark beetles disrupt the nutrient and water transport, gradually weakening and eventually killing the tree. Some bark beetles, e.g. European spruce bark beetle *Ips typographus*, often infest groups of trees, resulting in distinct and spatially-restricted patches of snags or dying trees. These habitat patches with higher tree mortality, in turn, open up gaps in the canopy, thereby initiating profound changes in the microclimatic and trophic conditions (Biedermann et al. 2019). Canopy gaps resulting from bark beetle activity increase light penetration, reduce canopy interception and enhance soil moisture, all of which accelerate organic matter decomposition and nutrient cycling (Siegert et al. 2024). Unlike the openings resulting from abiotic disturbances such as hurricane-force winds or avalanches, those produced by bark beetles are distinctive owing to presence of large numbers of snags (Bouget and Duelli 2004). However, trees within such gaps that manage to survive bark beetle outbreaks are also more susceptible to damage from wind, snow and falling adjacent trees (Palm-Hellenurm et al. 2024), potentially leading to multiple injuries such as bark scratches and branch or trunk breakage. All these factors create within a gap a habitat patch with diverse tree assemblages, including new tree species, numerous snags in various stages of decay, as well as living trees of different health and crown architecture, each potentially supporting particular TreMs. Consequently, a bark beetle-induced canopy gap may harbour a rich, abundant and diversified TreM assemblage. Since bark beetle-induced tree mortality is spatially restricted, the post-outbreak stands form a mosaic of disturbed patches surrounded by undisturbed, homogeneous closed-canopy forest (Müller et al. 2008), both probably characterized by distinctive TreM assemblages.

Bark beetle activity may mediate the biodiversity of temperate coniferous forests. Increased tree mortality from bark beetles influences the species composition and enhances bat activity (Piksa et al. 2022; Rachwald et al. 2022). Bark beetle-induced gaps host nearly 25% more bird species and 40% more individual birds than closed-canopy stands (Przepióra et al. 2020). Moreover, canopy gaps produced in this way give rise to altered species compositions and abundances of certain groups of insects (Tykarski and Knutelski 2010). As many species depend on TreMs, a plausible explanation for such a biodiversity boost is the rich and abundant assemblage of TreMs within bark beetle gaps. However, the impact of bark beetle activity on TreM assemblages has yet to be explored. In this study, we aimed (1) to assess the profile, i.e. the frequency of occurrence and density, of distinctive TreMs, as well as the richness, density and diversity of the TreM assemblage within an old-growth montane coniferous forest affected by bark beetle activity, and (2) to explore the potential effect of bark beetles on TreM numbers and composition turnover at the individual tree and forest stand scales. Canopy gaps resulting from bark beetle activity trigger an ecological cascade that alters tree species compositions, tree age structures, microclimates and deadwood resources, so we expected them to host a richer, more abundant and diverse TreM assemblage than a closed-canopy coniferous forest. We also anticipated changes in the abundance and frequency of specific TreMs within the disturbed patches. An understanding of the potential role of bark beetles as ecosystem engineers that mould TreM profiles is essential for evaluating the biodiversity dynamics of temperate coniferous forests.

## 2. Methods

### 2.1. Study area

The study was carried out in the Tatra Mountains (Mts.), which straddle the Polish-Slovakian border (Fig. 1a). They are protected by the Tatra National Park (TNP, established in 1954, covering 21,197 ha in the Polish part of the range) and the Tatranský Národný Park (established in 1949, covering 73,800 ha in the Slovakian part of the range; Fig. 1b). The Tatras are the highest range in the Carpathian Mountains and the only alpine-type mountains in Central Europe. Their highest peak reaches 2655 m above mean sea level (amsl), the average temperature ranges from –4°C on the mountain summits to 5°C at lower elevations, and there are from 135 to 230 days of continuous snow cover per year (Hess 1996).

**Fig. 1.**
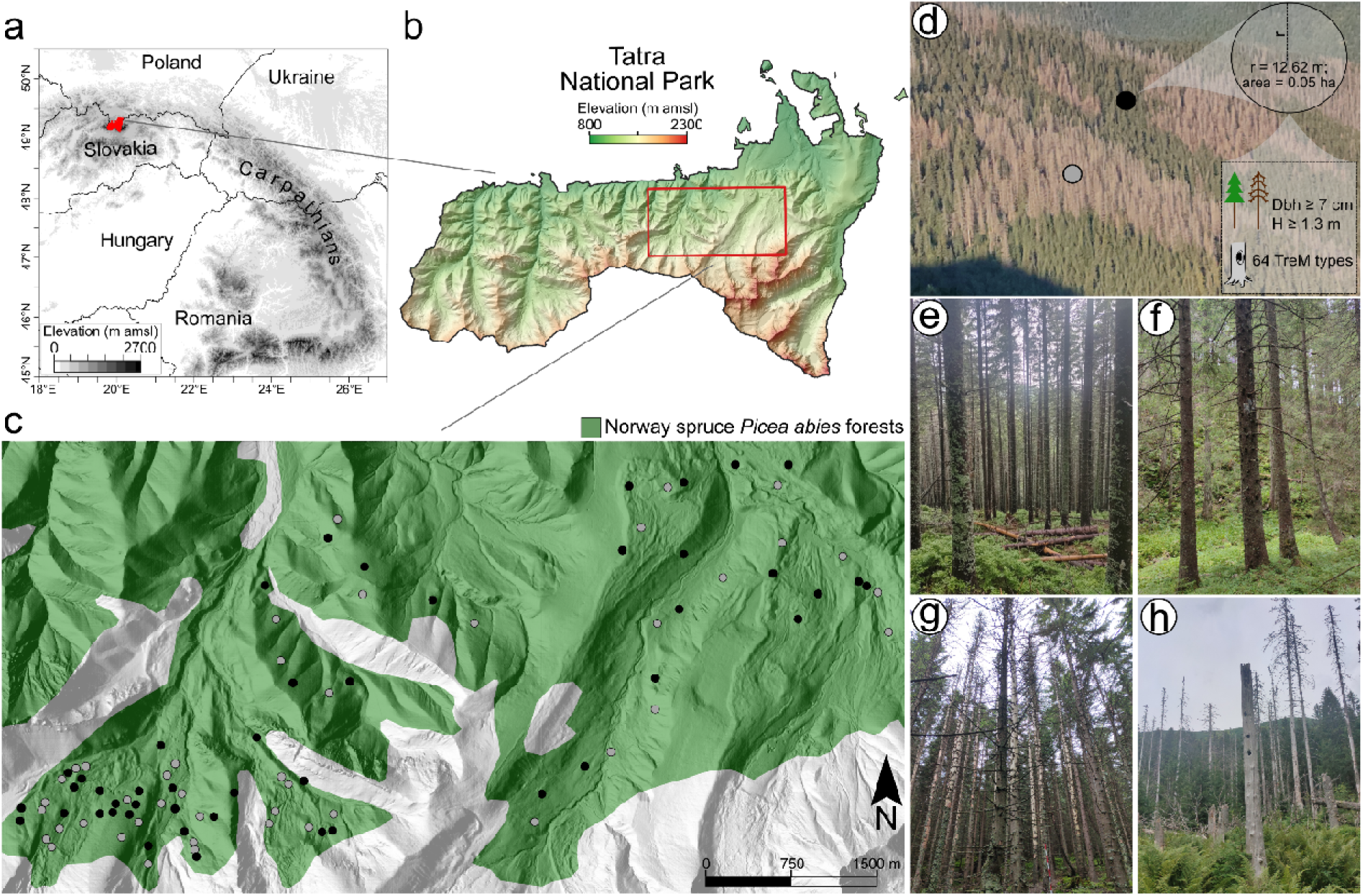
(**a**) Location of the study region – the Tatra National Park (TNP) covering the Polish part of the Tatra Mountains on the Polish-Slovakian border in the northern Carpathians, **(b)** location of the study area in relation to the topography and elevation of the TNP, **(c)** locations of the study plots (44 canopy gaps – grey circles and 46 reference sites – black circles) within the upper montane Norway spruce *Picea abies*-dominated old-growth forest, **(d)** protocol for tree measurements and Tree-related Microhabitat (TreM) cataloguing performed on each plot (Dbh – diameter at breast height (∼1.3 m); H – height of a tree; “TreM types” refers to the structures listed in Kraus et al. 2016), **(e)** closed-canopy Norway spruce-dominated forest with high or **(f)** low density of living trees that served as reference sites, and **(g)** bark beetle infestation patch with recently killed trees or **(h)** large infestation patches with dead standing trees in various stages of decomposition acting as canopy gaps (photos d-h by Fabian Przepióra).

Distinct climate-vegetation zones are discernible in the Tatra Mts. The lower montane zone, from 550 to 1200 m amsl, originally covered by lower montane forests with European beech *Fagus sylvatica* and silver fir *Abies alba*, have become transformed into Norway spruce *Picea abies* stands. The upper montane zone, naturally dominated by Norway spruce forests, extends up to the tree line at 1550 m amsl. Above the tree line, as far as 1800 m amsl, is the subalpine zone, dominated by dwarf pine *Pinus mugo* thickets, and higher up, all the way to the mountain summits, is the alpine zone with its characteristic grassland communities (Piękoś-Mirkowa and Mirek 1996). The majority of habitats within the climate-vegetation zones, comprising approximately 60% of the TNP’s total area, are subject to strict conservation (Havašová et al. 2017).

The study plots were located within the strictly protected old-growth forests in the upper montane zone (Fig. 1c), where Norway spruce is dominant with admixed rowan *Sorbus aucuparia*, the respective average volumes of the living trees of these species being 280 and 2 m^3^ ha^-1^. Within this zone, sycamore *Acer pseudoplatanus*, silver fir, European beech and European larch *Larix decidua* are also locally present; the volume of each of these species does not exceed 1 m^3^ ha^-1^ (Bodziarczyk et al. 2019). The spatial and temporal dynamics of these forests are governed by very strong Föhn-type winds, blowing mostly from the S and SW (Hess 1996), and also by the activities of cambiophages, particularly the European spruce bark beetle (Havašová et al. 2017). Windthrow events often trigger large-scale, natural renewal of Norway spruce, producing vast, single-layered stands of uniform age that continue to be vulnerable to environmental stressors such as wind, snow and/or insects (Holeksa and Szwagrzyk 2004). Larger and smaller patches of bark beetle infestation and windthrows currently cover about 30% of the Norway spruce forest area in the Tatra Mts. (Migas-Mazur et al. 2021).

The majority of the bark beetle infestation patches in the study area came into existence between 2012 and 2014 (Havašová et al. 2017; Przepióra et al. 2020). The strict conservation regulations applicable to this zone prohibit the removal of living trees or snags, salvage logging and tree planting. This non-interventionist (passive) management approach, coupled with the presence of multiple infestation patches, i.e. local concentrations of snags and fallen logs, has yielded an average deadwood volume of 248 m^3^ ha^-1^ in the upper montane spruce forests (Bodziarczyk et al. 2019). With their abundance of deadwood and diverse microclimatic and solar radiation conditions, bark beetle infestation patches in the TNP have been identified as diversity hotspots for vascular plants (Szwagrzyk et al. 2019), lichenized and lichenicolous fungi (Czarnota 2012), as well as birds, including three species of woodpecker, i.e. black woodpecker *Dryocopus martius*, great spotted woodpecker *Dendrocopos major* and Eurasian three-toed woodpecker *Picoides tridactylus* (Przepióra et al. 2020). In addition, the upper montane forests within the strictly protected zone are inhabited by large mammals such as brown bear *Ursus arctos* and red deer *Cervus elaphus*, which play pivotal roles in forest ecosystems by browsing or debarking living trees, foraging on saplings and dispersing seeds (Szwagrzyk et al. 2020; García-Rodríguez et al. 2021). These forests also host large saproxylic invertebrates like the sawyer beetle *Monochamus sartor*, which bores exit holes up to 1 cm in diameter (Tykarski and Knutelski 2010).

### 2.2. Field methods

The survey was carried out on 90 circular study plots 12.62 m in radius (0.05 ha) (Fig. 1c-d), located within bark beetle infestation patches defined as areas surrounded by living, closed-canopy tree stands, in which the feeding of cambiophagous insects had caused small-scale die-back of trees leading to gaps in the canopy (forest with canopy closure < 50%; henceforth: canopy gaps; Fig. 1g-h). The reference plots were located in the closed-canopy forests surrounding the canopy gaps (henceforth: reference sites; Fig. 1e-f). The study was so designed as to enable investigation of the connection between bark beetle activity and TreMs and used a ∼1:1 gap-to-reference forest ratio, unlike the actual proportions of these habitat stages in the Tatra Mts., which is currently ∼3:7 (Migas-Mazur et al. 2021).

The canopy gaps were delimited around the borders of bark beetle infestation patches (N = 44) chosen at random from a pool of all the bark beetle infestation patches visualized on an orthophotomap of the surveyed area. The reference plots were then designated at random in the surrounding closed-canopy forest, ensuring a minimum distance of 50 m from the plots in the canopy gaps (N = 46, Fig. 1d) and a minimum distance of 50 m between adjacent plots, i.e. more than twice the average stand height in these forests (Przepióra et al. 2020). The canopy gaps included infestation patches of various sizes, from relatively small openings to areas where die-back had affected several ha of forest; the median area of an infestation patch was thus 0.53 ha (quartile range 0.16–2.40 ha, range 0.05–20.58 ha). The mean distance between the ultimately selected adjoining canopy gaps and reference sites was 209.6 m ±97.4 SD (range 62.8–408.5 m).

The fieldwork was carried out between August and September in 2020-2023. On each plot, every living tree or snag with a diameter measured at a height of 1.3 m above the ground (hereafter, diameter) ≥ 7 cm was examined for the presence of TreMs (Fig. 1d). The species and living status of each tree (living tree *vs*. snag) were identified and its diameter measured. TreMs were catalogued in accordance with the typology for temperate forests (Kraus et al. 2016) in order to confirm the presence or absence of a structure on a given tree (Przepióra and Ciach 2022). The observer effect was eliminated in that the TreMs were catalogued by one person only (the first author of this paper). Each tree was scrutinized for at least 3 minutes, with the upper parts of the trunk and crown being examined using binoculars.

### 2.3. Remotely-sensed data

The topographic features were derived using open-source Light Detection and Ranging (LiDAR) data, collected via airborne laser scanning with a density of 4 points m^-2^ (https://www.geoportal.gov.pl/en/data/lidar-measurements-lidar). The data was compiled into a multi-layer raster with a resolution of 22.36 m (approximately 500 m^2^, matching the study plot size). Prior to computing the topographic features, the raw LiDAR data was converted into a Digital Elevation Model (DEM). Features calculated for each raster cell included elevation, slope inclination, Topographic Position Index (TPI), northness and eastness. Elevation, representing the height of a pixel relative to mean sea level (m amsl), and slope inclination (degrees) were derived from the DEM. Terrain ruggedness, indicated by TPI, was determined as the elevation difference between a central pixel and the average elevation of its eight neighbouring cells. Large negative TPI values denoted valleys or gully bottoms, large positive values indicated ridges, hilltops or mountain summits, and near-zero values signified either flat areas (with a near-zero slope) or mountain slopes (with a non-zero slope) (Weiss 2001). Northness was calculated as the cosine of the aspect, ranging from 1 (due north) to –1 (due south), with 0 indicating no north or south aspect. Eastness was computed as the sine of the aspect, ranging from 1 (due east) to –1 (due west), with 0 indicating no east or west aspect.

### 2.4. Data handling and analyses

#### 2.4.1. Tree traits and TreM richness at the individual tree level

A total of 4052 trees – 2512 living ones and 1540 snags – were examined during the fieldwork. As 99.2% were Norway spruces, all the analyses at the individual tree level were performed for that species (2482 living trees and 1537 snags; Table S1). The basal area (the cross-sectional area of the tree trunk at breast height expressed in m^2^ ha^-1^) and TreM richness (expressed as the total number of TreM types recorded on an individual tree) of each living tree and snag were calculated. The topographic characteristics, including elevation, slope inclination, TPI, northness and eastness, were read for each individual living tree and snag from the cell of the multi-layer raster (see Remotely-sensed data section) where the tree/snag was located.

The mean, standard deviation (SD) and range of TreM richness and diameter were calculated for the following groups: (1) living trees and snags pooled, (2) living trees and snags separately, and (3) living trees and snags located separately in canopy gaps or reference sites. Student’s t-test was used to test the differences in mean TreM richness and mean diameter between living trees and snags, regardless of location, as well as between living trees and snags located in either canopy gaps or reference sites.

Prior to the modelling procedures, the explanatory variables were tested for collinearity using Pearson’s correlation. Variable pairs with a correlation coefficient r ≤ 0.6 were included in the analyses (Fig. S1a-b). The relationships between the individual characteristics and TreM richness found on a given tree located in a canopy gap or reference site were analysed using Generalized Linear Mixed Models (GLMMs) with a Poisson error distribution and log link function. First, the models with the identification number of the study plot included as a random effect were built using all possible combinations of six continuous explanatory variables **–** diameter, elevation, slope inclination, TPI, northness and eastness **–** and the categorical variable of living status (living tree *vs*. snag). Then, the model with the lowest Akaike Information Criterion corrected for small sample size (AICc) was selected as the best model for trees located in canopy gaps and reference sites. The AICc of the best model for trees located in canopy gaps with six fixed effects was not different (ΔAICc = 1.3), while the AICc of the best model for trees located in reference sites with seven fixed effects and random effect (SD = 0.39) was lower than that of the model with all explanatory variables (ΔAICc = 302.9). The residuals of the best model for trees located in canopy gaps (dispersion = 0.76; p < 0.001; simulation-based dispersion tests) and reference sites (dispersion = 0.68; p < 0.001; simulation-based dispersion tests) exhibited dispersion. The zero-inflation tests showed no significant zero-inflation for trees located in canopy gaps (ratioObsSim = 0.93; p = 0.304) and reference sites (ratioObsSim = 0.89; p = 0.064). Pairwise comparison of TreM richness between canopy gaps and reference sites, predicted for living trees or snags, was conducted using a two-sample t-test for dependent samples at defined tree diameter thresholds, measured at 10 cm-width intervals (i.e. 10 cm, 20 cm, 30 cm, …, 70 cm). For each threshold, the difference and the standard error of the difference between model-predicted values for canopy gaps and reference sites were used to compute a t-statistic and a p-value, in order to assess whether the difference was other than zero.

#### 2.4.2. Habitat characteristics and TreM indices at the study plot level

All the analyses at the study plot level were based on the whole dataset, i.e. 4052 trees (Table S1). A set of characteristics describing the forest stand structure and composition, topography and TreM assemblage was calculated for each plot, and the mean, SD and range of these characteristics were calculated for all 90 study plots and separately for 44 canopy gaps and 46 reference sites. The forest stand structure characteristics included the density and basal area calculated separately for living trees and snags, while the forest stand composition characteristic was the number of tree species calculated on the basis of living trees and snags pooled. The topographic characteristics, i.e. elevation, slope inclination, TPI, northness and eastness, were read from the cell of the multi-layer raster (see Remotely-sensed data section), where the centres of the study plots were located.

The characteristics of the TreM assemblage were calculated on the basis of both living trees and snags pooled and included three TreM indices: TreM richness, i.e. the total number of TreM types recorded on a study plot; TreM density, i.e. the total density of trees bearing TreMs (if a tree bore several TreM types, it was counted once for each TreM type) extrapolated to 1 ha (expressed as TreM-bearing trees ha^-1^) (Paillet et al. 2017); TreM diversity, i.e. the Shannon-Wiener diversity index calculated on the basis of TreM types recorded and their relative abundance (the density of trees bearing a given TreM type in the density of all TreM-bearing trees recorded on a study plot) (Przepióra and Ciach 2022).

In accordance with the typology of Kraus et al. (2016), we grouped all TreM types into twenty TreM groups and eight TreM forms. We calculated the density of TreM-bearing trees, i.e. the number of trees bearing a given TreM extrapolated to 1 ha, and the frequency, defined as the percentage of study plots with a given TreM, for each group and form. We also calculated the mean, SD and range of the density of TreM-bearing trees and the frequency, defined as the number of plots with a given TreM, for each type, separately for canopy gaps and reference sites. If particular TreM groups/forms were represented by several TreM types/groups on one tree, we counted the presence of a group/form only once. Therefore, the calculations of density and frequency of given TreM groups or forms on an individual tree were based on the occurrence of at least one TreM type from a given group or form, respectively. The differences in characteristics describing the forest stand structure and composition, topography and TreM indices between canopy gaps and reference sites were tested using Student’s t-test. The differences in density of TreM-bearing trees or frequency of a given TreM type, group or form between canopy gaps and reference sites were tested using the Mann-Whitney U-test and the chi-square goodness-of-fit test with Yates’ correction, respectively.

The relationships between habitat characteristics and the three TreM indices found in a given study plot were analysed using GLMMs with a Poisson error distribution and log link function for TreM richness, with a Gaussian error distribution and identity link function for TreM density, and with a Gamma error distribution and log link function for TreM diversity. Models were built separately for canopy gaps and reference sites. The best models were selected using the same procedure as with the model at the individual tree level, based on a set of eight continuous explanatory variables **–** number of tree species, density of living trees, density of snags, elevation, slope inclination, TPI, northness and eastness. The AICc of each best model was lower than that of the corresponding model with all the explanatory variables: TreM richness models for canopy gaps (ΔAICc = 7.2) and reference sites (ΔAICc = 6.4), TreM density models for canopy gaps (ΔAICc = 5.9) and reference sites (ΔAICc = 4.0), and TreM diversity models for canopy gaps (ΔAICc = 3.9) and reference sites (ΔAICc = 6.9). Since in none of the models did the residuals of the final models exhibit overdispersion (p = 0.200 – 0.960) or zero-inflation (p = 1.000), random effects were not added to any models, i.e. not for TreM richness or TreM density or TreM diversity, regardless of whether they concerned canopy gaps or reference sites.

The relationships between habitat characteristics and the density of trees bearing specific TreM types in the study plots were analysed using Redundancy Analysis (RDA). RDA extends principal component analysis by constraining results, i.e. study plot and TreM type scores with explanatory variables. Prior to the analyses, the response variable (density of TreM-bearing trees) was transformed using Hellinger’s square-root transformation. A set of eight continuous explanatory variables **–** number of tree species, density of living trees, density of snags, elevation, slope inclination, TPI, northness and eastness **–** was initially included in the global RDA. To reduce the number of variables, permutational multivariate analysis of variance (PERMANOVA) was performed with all possible combinations, with the set of variables with the lowest AIC being selected. The final RDA was characterized by a ΔAIC of 2.1 compared to the global RDA.

All the analyses were conducted in R 3.5.0 (CoreTeam R. 2017). Topographic variables were computed using the terra package (Hijmans et al. 2022). GLMMs were performed with the glmmTMB package (Magnusson et al. 2017), model diagnostics were done using DHARMa package (Hartig and Hartig 2017) and the vegan package (Dixon 2003) was used for RDA.

## 3. Results

### 3.1. TreM richness at the individual tree level

The mean TreM richness found on living trees and snags pooled was 1.8 ±1.8 SD TreM types per tree. The mean TreM richness found on living trees was 1.1 ±1.2 SD TreM types per tree and was lower (t = 32.7; p < 0.001) than the mean TreM richness found on snags (3.0 ±2.0 SD TreM types per tree). Although snags in canopy gaps had a higher TreM richness than those in reference sites, the TreM richness of living trees did not differ significantly between the two groups of study plots (Table 1a).

**Table 1.** (**a**) Mean ±SD and range of diameter at breast height of living trees or snags and Tree-related Microhabitat (TreM) richness (total number of TreM types recorded on an individual tree) found on living trees or snags, calculated separately for trees located in canopy gaps and reference sites; **(b)** density of living trees, density of snags, basal area of living trees, basal area of snags, number of tree species; **(c)** elevation, slope inclination, Topographic Position Index (TPI), northness, eastness; **(d)** TreM richness (total number of TreM types recorded on a study plot), TreM density (density of TreM-bearing trees ha^−1^), and TreM diversity (Shannon-Wiener index), calculated on the basis of living trees and snags pooled, separately for canopy gaps and reference sites, located in Norway spruce *Picea abies*-dominated old-growth forests in the Tatra Mountains (Poland).

| Characteristics | Unit | Canopy gaps |  |  | Reference sites |  |  | t | p |
| --- | --- | --- | --- | --- | --- | --- | --- | --- | --- |
|  |  | Mean | SD | Range | Mean | SD | Range |  |  |
| **(a) Individual tree traits** |  |  |  |  |  |  |  |  |  |
| Diameter at breast height of living trees | cm | 21.6 | 9.1 | 7.0-57.5 | 22.6 | 9.8 | 7.0-75.4 | -1.76 | 0.079 |
| Diameter at breast height of snags | cm | 29.3 | 12.0 | 7.0-68.0 | 15.1 | 7.6 | 7.0-62.5 | 28.09 | 0.000 |
| TreM richness of living trees | TreM types per tree | 1.2 | 1.2 | 0.0-7.0 | 1.1 | 1.2 | 0.0-7.0 | 1.76 | 0.080 |
| TreM richness of snags | TreM types per tree | 3.5 | 2.0 | 0.0-14.0 | 2.0 | 1.7 | 0.0-10.0 | 14.56 | 0.000 |
| **(b) Stand structure** |  |  |  |  |  |  |  |  |  |
| Density of living trees | trees·ha<sup>-1</sup> | 145.9 | 170.4 | 0.0-800.0 | 952.6 | 363.1 | 320.0-1800.0 | -13.59 | 0.000 |
| Density of snags | trees·ha<sup>-1</sup> | 467.7 | 202.4 | 100.0-1080.0 | 222.2 | 180.2 | 0.0-780.0 | 6.03 | 0.000 |
| Basal area of living trees | m<sup>2</sup>·ha<sup>-1</sup> | 6.3 | 8.9 | 0.0-44.3 | 45.9 | 8.3 | 27.9-61.8 | -21.81 | 0.000 |
| Basal area of snags | m<sup>2</sup>·ha<sup>-1</sup> | 37.0 | 12.6 | 13.9-64.2 | 5.0 | 4.5 | 0.0-23.8 | 15.90 | 0.000 |
| Number of tree species | N | 1.1 | 0.3 | 1.0-2.0 | 1.3 | 0.6 | 1.0-4.0 | -1.44 | 0.155 |
| **(c) Topography** |  |  |  |  |  |  |  |  |  |
| Elevation | m amsl | 1336.7 | 76.7 | 1140.0-1457.0 | 1316.0 | 79.9 | 1095.0-1454.0 | 1.26 | 0.212 |
| Slope inclination | degrees | 15.3 | 9.0 | 1.5-37.8 | 13.6 | 8.2 | 1.6-33.1 | 0.95 | 0.343 |
| TPI | Δm | 0.4 | 1.9 | -3.8-6.8 | -0.1 | 2.7 | -9.3-5.8 | 1.01 | 0.317 |
| Northness | value from -1 to 1 | 0.7 | 0.5 | -0.9-1.0 | 0.6 | 0.5 | -0.8-1.0 | 0.90 | 0.373 |
| Eastness | value from -1 to 1 | 0.0 | 0.6 | -1.0-1.0 | 0.1 | 0.6 | -0.1-1.0 | -0.57 | 0.570 |
| **(d) TreM indices** |  |  |  |  |  |  |  |  |  |
| TreM richness | TreM types per plot | 12.3 | 3.9 | 6.0-23.0 | 13.5 | 3.6 | 6.0-21.0 | -1.51 | 0.134 |
| TreM density | TreM-bearing trees·ha<sup>-1</sup> | 1796.8 | 758.1 | 600.0-3160.0 | 1490.0 | 754.8 | 380.0-3980.0 | 1.92 | 0.058 |
| TreM diversity | Shannon-Wiener index | 0.9 | 0.1 | 0.6-1.1 | 0.8 | 0.2 | 0.4-1.3 | 1.36 | 0.178 |

The TreM richness of an individual tree was correlated with its living status, being higher for snags in both canopy gaps and reference sites (Table 2a). TreM richness increased with diameter and elevation for trees in both canopy gaps and reference sites, and with eastness for trees in canopy gaps (Table 2a). TreM richness decreased with TPI for trees in canopy gaps and reference sites, and with northness for trees in canopy gaps (Table 2a). Living trees with diameters of 10 cm, 20 cm and 30 cm (Fig. 2a) and snags with diameters of 10 cm and 20 cm (Fig. 2b) had a higher TreM richness in canopy gaps, while living trees with diameters of 60 cm and 70 cm (Fig. 2a), and snags with diameters of 50 cm, 60 cm and 70 cm (Fig. 2b) had a higher TreM richness in reference sites.

**Fig. 2.**
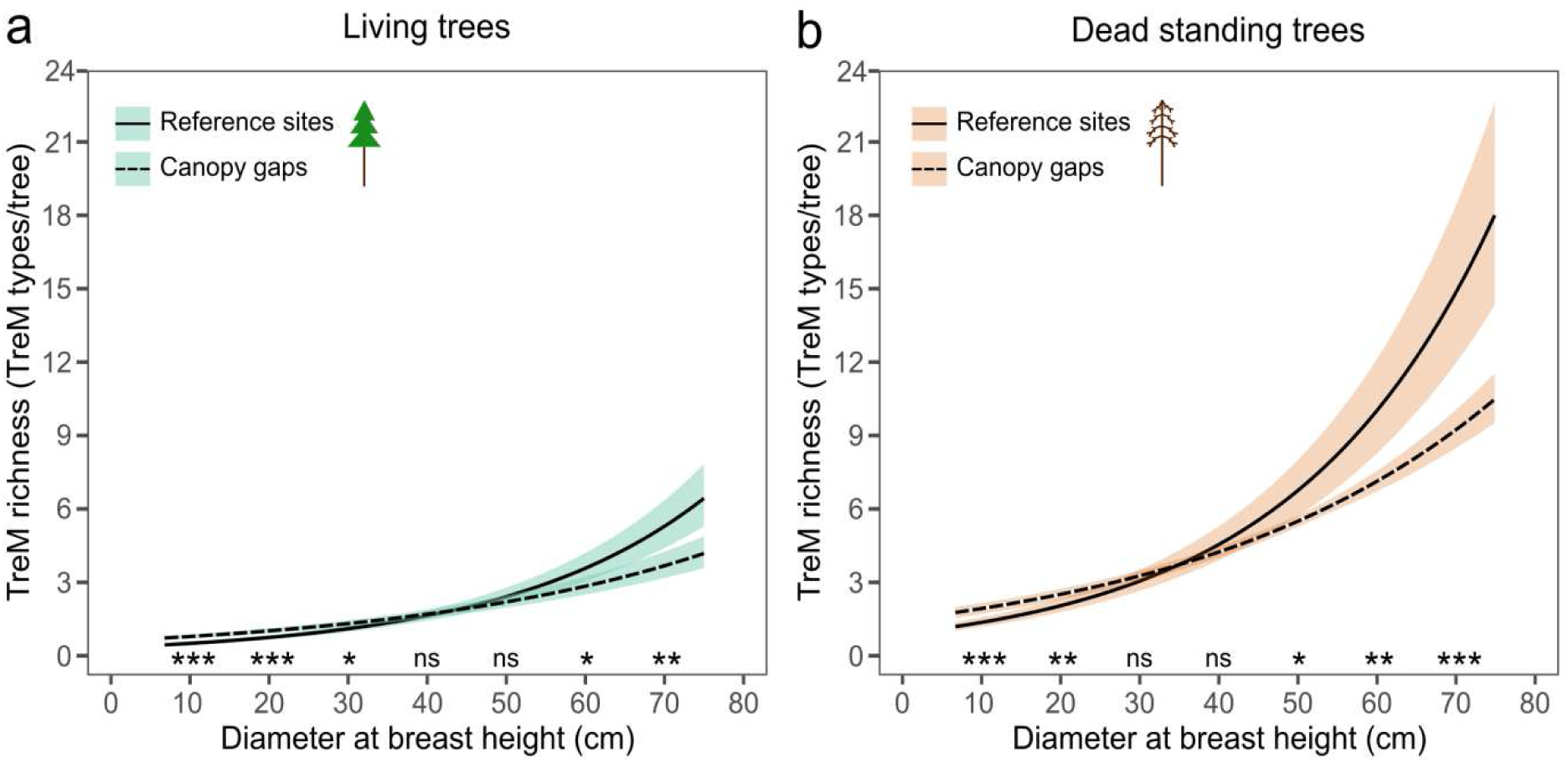
The relationships between Tree-related Microhabitat (TreM) richness (the total number of TreM types recorded on an individual tree) and diameter at breast height of **(a)** living trees (turquoise ribbons) and **(b)** dead standing trees (copper ribbons) located in canopy gaps (dashed lines) and reference sites (solid lines) in Norway spruce *Picea abies*-dominated old-growth forests in the Tatra Mountains (Poland). Mean (solid or dashed lines) and confidence intervals (ribbons) are a product of generalized linear-mixed models (Table 2a). Significance levels for the pairwise comparison of model predictions at specific diameter thresholds are indicated as follows: ***p ≤ 0.001, **p ≤ 0.01, *p ≤ 0.05, and ns for non-significant comparisons.

**Table 2.** Models describing the relationships between. **(a)** Tree-related Microhabitat (TreM) richness at the individual tree level (the total number of TreM types recorded on an individual tree) and fixed effects: living status (living *vs*. dead standing tree), diameter at breast height, elevation, slope inclination, Topographic Position Index (TPI), northness and eastness, calculated separately for trees located in canopy gaps (N = 44) and reference sites (N = 46); **(b)** TreM richness at the plot level (total number of TreM types recorded on a study plot); **(c)** TreM density (density of TreM-bearing trees ha^-1^); **(d)** TreM diversity (Shannon-Wiener index) recorded in canopy gaps (N = 44) and reference sites (N = 46) in the Norway spruce *Picea abies*-dominated old-growth forest in the Tatra Mountains (Poland), and habitat characteristics (fixed effects): number of tree species, density of living trees, density of snags (dead standing trees), basal area of living trees, basal area of snags, elevation, slope inclination and TPI.

| Model | Canopy gap |  |  |  | Reference site |  |  |  |
| --- | --- | --- | --- | --- | --- | --- | --- | --- |
|  | Estimate | SE | z | Pr(>\|z\|) | Estimate | SE | z | Pr(>\|z\|) |
| **(a)** Tree microhabitat richness (individual tree level) |  |  |  |  |  |  |  |  |
| Intercept | -2.455 | 0.31 | -7.98 | 0.000 | -5.272 | 1.14 | -4.61 | 0.000 |
| Living status (dead tree) | 0.926 | 0.06 | 16.15 | 0.000 | 1.034 | 0.04 | 23.59 | 0.000 |
| Diameter at breast height | 0.026 | 0.00 | 20.14 | 0.000 | 0.039 | 0.00 | 23.53 | 0.000 |
| Elevation | 0.002 | 0.00 | 6.50 | 0.000 | 0.003 | 0.00 | 3.77 | 0.000 |
| Slope inclination |  |  |  |  | -0.015 | 0.01 | -1.87 | 0.061 |
| TPI | -0.029 | 0.01 | -2.99 | 0.003 | -0.047 | 0.02 | -1.97 | 0.049 |
| Northness | -0.086 | 0.03 | -2.51 | 0.012 | -0.075 | 0.12 | -0.61 | 0.543 |
| Eastness | 0.097 | 0.03 | 3.28 | 0.001 | 0.002 | 0.10 | 0.02 | 0.983 |
| **(b)** Tree microhabitat richness (plot level) |  |  |  |  |  |  |  |  |
| Intercept | 0.470 | 0.81 | 0.58 | 0.560 | 0.881 | 0.71 | 1.24 | 0.216 |
| Density of living trees | 0.001 | 0.00 | 4.08 | 0.000 | -0.000 | 0.00 | -1.73 | 0.084 |
| Density of snags |  |  |  |  | 0.000 | 0.00 | 1.91 | 0.056 |
| Number of tree species |  |  |  |  | 0.124 | 0.07 | 1.86 | 0.063 |
| Elevation | 0.001 | 0.00 | 2.38 | 0.017 | 0.001 | 0.00 | 2.42 | 0.016 |
| **(c)** Tree microhabitat density (plot level) |  |  |  |  |  |  |  |  |
| Intercept | 0.967 | 1239.58 | 0.00 | 0.999 | -1.487 | 1183.64 | 0.00 | 0.999 |
| Density of living trees | 0.208 | 0.14 | 1.50 | 0.134 |  |  |  |  |
| Density of snags | 2.045 | 0.30 | 6.80 | 0.000 | 1.818 | 0.28 | 6.58 | 0.000 |
| Elevation |  |  |  |  | 0.908 | 0.89 | 1.02 | 0.308 |
| Slope inclination |  |  |  |  | -11.825 | 7.36 | -1.61 | 0.108 |
| TPI |  |  |  |  | -42.804 | 27.42 | -1.56 | 0.119 |
| **(d) Tree microhabitat diversity (plot level)** |  |  |  |  |  |  |  |  |
| Intercept | 2.699 | 0.42 | 6.37 | 0.000 | 1.169 | 0.10 | 11.35 | 0.000 |
| Density of living trees | -0.000 | 0.00 | -2.54 | 0.011 | 0.000 | 0.00 | 2.38 | 0.017 |
| Density of snags | 0.000 | 0.00 | 1.86 | 0.063 |  |  |  |  |
| Number of tree species |  |  |  |  | -0.142 | 0.05 | -3.04 | 0.002 |
| Elevation | -0.001 | 0.00 | -3.92 | 0.000 |  |  |  |  |

### 3.2. Habitat traits and TreM indices at the plot level

Seven tree species were found in the Norway spruce-dominated forest (Table S1), and the mean number of species in canopy gaps and reference sites did not differ (Table 1b). The mean density and mean basal area of living trees were lower in canopy gaps than in reference sites, whereas the mean density and mean basal area of snags were higher in canopy gaps than in reference sites (Table 1b). The study plots were situated on slopes with a mean inclination of 14.4 ±8.6 SD degrees, at a mean elevation of 1326.0 ±78.6 SD m amsl and a mean northness of 0.6 ±0.5 SD. None of the topographic parameters, i.e. elevation, slope inclination, TPI, northness or eastness, differed between canopy gaps and reference sites (Table 1c). The plots were characterized by a mean TreM richness of 12.9 ±3.7 SD TreM types per plot (range 6.0–23.0), a mean TreM density of 1640.0 ±767.8 SD TreM-bearing trees ha^-1^ (range 380.0–3980.0) and a mean TreM diversity of 0.8 ±0.2 SD (range 0.4–1.3). None of the three TreM indices, i.e. richness, density or diversity of TreMs, differed between canopy gaps and reference sites, although the higher TreM density in canopy gaps did approach significance (Table 1d).

### 3.3. TreM assemblage in canopy gaps and reference sites

A total of 8 forms (Table S2a), 19 groups (Table S2b) and 49 types (Table 3) of TreMs were identified in the studied Norway spruce-dominated forest. The forms with the highest mean density in canopy gaps were epiphytes and bark, while the forms with the highest mean density in reference sites were epiphytes and deformations / growth forms (Table S2a). The groups with the highest mean density in canopy gaps were epiphytic crypto– and phanerogams, bark pockets, fungal fruiting bodies, insect galleries and bore holes, while the groups with the highest mean density in reference sites were epiphytic crypto– and phanerogams, root buttress cavities and bark pockets (Table S2b).

**Table 3.** Mean, SD and range of Tree-related Microhabitat (TreM) densities (density of trees bearing a given TreM ha^-1^) and frequencies (number of plots with a given TreM) of types, found in canopy gaps (N = 44) and reference sites (N = 46) in the Norway spruce *Picea abies*-dominated old-growth forests in the Tatra Mountains (Poland). Significant results (p < 0.05) are shown in bold. The codes of TreM types are adapted from Kraus et al. (2016). The full names and descriptions of TreM types are given in Table S2.

| Code | Density |  |  |  |  |  | Frequency |  |  |  |  |  |
| --- | --- | --- | --- | --- | --- | --- | --- | --- | --- | --- | --- | --- |
|  | Canopy gaps |  |  | Reference sites |  |  | z | p | Canopy gaps | Reference sites | χ<sup>2</sup> | p |
|  | Mean | SD | Range | Mean | SD | Range |  |  |  |  |  |  |
| CV11 | 0.5 | 3.0 | 0.0-20.0 | 0.0 | 0.0 | 0.0-0.0 | -1.00 | 0.317 | 1 | 0 | 0.00 | 0.982 |
| **CV12** | **7.7** | **15.1** | **0.0-60.0** | **1.3** | **5.0** | **0.0-20.0** | **-2.67** | **0.008** | **12** | **3** | **5.56** | **0.018** |
| CV13 | 0.5 | 3.0 | 0.0-20.0 | 0.0 | 0.0 | 0.0-0.0 | -1.00 | 0.317 | 1 | 0 | 0.00 | 0.982 |
| CV14 | 9.5 | 17.0 | 0.0-60.0 | 10.9 | 15.0 | 0.0-60.0 | -0.88 | 0.381 | 13 | 19 | 0.89 | 0.345 |
| CV15 | 1.4 | 5.1 | 0.0-20.0 | 0.4 | 2.9 | 0.0-20.0 | -1.05 | 0.293 | 3 | 1 | 0.31 | 0.577 |
| CV21 | 2.3 | 6.4 | 0.0-20.0 | 2.2 | 9.6 | 0.0-60.0 | -0.75 | 0.452 | 5 | 3 | 0.19 | 0.663 |
| CV22 | 0.5 | 3.0 | 0.0-20.0 | 2.6 | 8.0 | 0.0-40.0 | -1.62 | 0.104 | 1 | 5 | 1.47 | 0.226 |
| CV23 | 0.9 | 4.2 | 0.0-20.0 | 0.0 | 0.0 | 0.0-0.0 | -1.44 | 0.150 | 2 | 0 | 0.56 | 0.455 |
| CV25 | 0.9 | 4.2 | 0.0-20.0 | 0.4 | 2.9 | 0.0-20.0 | -0.61 | 0.542 | 2 | 1 | 0.00 | 0.969 |
| CV26 | 0.9 | 4.2 | 0.0-20.0 | 0.4 | 2.9 | 0.0-20.0 | -0.61 | 0.542 | 2 | 1 | 0.00 | 0.969 |
| CV31 | 0.5 | 3.0 | 0.0-20.0 | 1.3 | 6.5 | 0.0-40.0 | -0.55 | 0.586 | 1 | 2 | 0.00 | 1.000 |
| CV44 | 1.8 | 5.8 | 0.0-20.0 | 1.3 | 5.0 | 0.0-20.0 | -0.44 | 0.657 | 4 | 3 | 0.00 | 0.951 |
| **CV51** | **149.1** | **148.1** | **0.0-520.0** | **57.0** | **92.4** | **0.0-540.0** | **-2.58** | **0.010** | **31** | **31** | **0.01** | **0.931** |
| CV52 | 0.9 | 6.0 | 0.0-40.0 | 0.9 | 4.1 | 0.0-20.0 | -0.51 | 0.613 | 1 | 2 | 0.00 | 1.000 |
| IN11 | 0.5 | 3.0 | 0.0-20.0 | 1.3 | 5.0 | 0.0-20.0 | -0.96 | 0.337 | 1 | 3 | 0.22 | 0.641 |
| IN12 | 0.9 | 4.2 | 0.0-20.0 | 1.3 | 5.0 | 0.0-20.0 | -0.40 | 0.692 | 2 | 3 | 0.00 | 1.000 |
| **IN13** | **2.7** | **9.2** | **0.0-40.0** | **13.5** | **20.7** | **0.0-100.0** | **-3.71** | **0.000** | **4** | **21** | **13.22** | **0.000** |
| **IN14** | **1.4** | **5.1** | **0.0-20.0** | **7.8** | **17.1** | **0.0-80.0** | **-2.11** | **0.035** | **3** | **10** | **2.93** | **0.087** |
| IN21 | 0.5 | 3.0 | 0.0-20.0 | 4.3 | 16.3 | 0.0-100.0 | -1.63 | 0.102 | 1 | 5 | 1.47 | 0.226 |
| IN23 | 0.0 | 0.0 | 0.0-0.0 | 0.4 | 2.9 | 0.0-20.0 | -0.96 | 0.339 | 0 | 1 | 0.00 | 1.000 |
| **IN24** | **7.7** | **15.7** | **0.0-60.0** | **0.4** | **2.9** | **0.0-20.0** | **-2.99** | **0.003** | **10** | **1** | **7.04** | **0.008** |
| **IN31** | **61.8** | **66.2** | **0.0-260.0** | **13.0** | **19.0** | **0.0-60.0** | **-3.91** | **0.000** | **31** | **19** | **6.60** | **0.010** |
| IN32 | 4.5 | 11.3 | 0.0-60.0 | 2.6 | 10.0 | 0.0-60.0 | -1.28 | 0.201 | 8 | 4 | 1.03 | 0.311 |
| DE11 | 0.5 | 3.0 | 0.0-20.0 | 0.0 | 0.0 | 0.0-0.0 | -1.00 | 0.317 | 1 | 0 | 0.00 | 0.982 |
| DE12 | 0.5 | 3.0 | 0.0-20.0 | 0.0 | 0.0 | 0.0-0.0 | -1.00 | 0.317 | 1 | 0 | 0.00 | 0.982 |
| **DE13** | **1.8** | **5.8** | **0.0-20.0** | **10.4** | **16.7** | **0.0-60.0** | **-3.02** | **0.003** | **4** | **16** | **7.17** | **0.007** |
| DE14 | 0.5 | 3.0 | 0.0-20.0 | 0.4 | 2.9 | 0.0-20.0 | -0.02 | 0.987 | 1 | 1 | 0.00 | 1.000 |
| **DE15** | **1.8** | **7.2** | **0.0-40.0** | **16.1** | **20.9** | **0.0-80.0** | **-4.31** | **0.000** | **3** | **22** | **16.86** | **0.000** |
| **BA11** | **312.3** | **142.1** | **80.0-660.0** | **93.0** | **99.4** | **0.0-480.0** | **-6.84** | **0.000** | **44** | **40** | **4.23** | **0.040** |
| **BA12** | **320.5** | **143.9** | **80.0-680.0** | **101.3** | **105.4** | **0.0-540.0** | **-6.73** | **0.000** | **44** | **40** | **4.23** | **0.040** |
| **BA21** | **9.1** | **18.5** | **0.0-80.0** | **24.3** | **28.6** | **0.0-100.0** | **-3.07** | **0.002** | **11** | **26** | **7.97** | **0.005** |
| **GR11** | **60.9** | **80.0** | **0.0-260.0** | **113.0** | **121.1** | **0.0-440.0** | **-2.47** | **0.014** | **23** | **36** | **5.62** | **0.018** |
| GR12 | 74.5 | 81.7 | 0.0-260.0 | 100.4 | 93.0 | 0.0-340.0 | -1.48 | 0.140 | 31 | 39 | 1.91 | 0.167 |
| GR13 | 14.1 | 23.1 | 0.0-60.0 | 25.2 | 46.7 | 0.0-180.0 | -0.66 | 0.507 | 14 | 17 | 0.08 | 0.771 |
| GR31 | 1.8 | 5.8 | 0.0-20.0 | 3.5 | 9.7 | 0.0-40.0 | -0.64 | 0.519 | 4 | 6 | 0.07 | 0.794 |
| **GR32** | **1.4** | **6.7** | **0.0-40.0** | **5.2** | **10.7** | **0.0-40.0** | **-2.33** | **0.020** | **2** | **10** | **4.36** | **0.037** |
| EP11 | 15.0 | 20.7 | 0.0-80.0 | 20.0 | 24.6 | 0.0-100.0 | -0.94 | 0.345 | 21 | 25 | 0.17 | 0.677 |
| **EP12** | **136.8** | **84.1** | **0.0-340.0** | **14.8** | **22.1** | **0.0-100.0** | **-7.60** | **0.000** | **43** | **20** | **28.99** | **0.000** |
| EP13 | 12.7 | 33.2 | 0.0-140.0 | 6.5 | 12.0 | 0.0-40.0 | -0.32 | 0.747 | 9 | 12 | 0.15 | 0.702 |
| EP14 | 2.7 | 18.1 | 0.0-120.0 | 2.2 | 7.6 | 0.0-40.0 | -1.27 | 0.204 | 1 | 4 | 0.76 | 0.385 |
| EP21 | 6.4 | 14.8 | 0.0-60.0 | 3.0 | 13.3 | 0.0-80.0 | -1.85 | 0.064 | 9 | 3 | 2.67 | 0.102 |
| EP31 | 80.9 | 100.4 | 0.0-360.0 | 103.5 | 160.5 | 0.0-640.0 | -0.20 | 0.841 | 33 | 33 | 0.01 | 0.911 |
| **EP32** | **430.0** | **246.8** | **40.0-1020.0** | **604.8** | **409.5** | **0.0-1700.0** | **-2.06** | **0.040** | **44** | **45** | **0.00** | **1.000** |
| **EP34** | **32.7** | **39.8** | **0.0-160.0** | **3.0** | **11.1** | **0.0-60.0** | **-5.37** | **0.000** | **28** | **4** | **27.28** | **0.000** |
| NE12 | 0.0 | 0.0 | 0.0-0.0 | 0.4 | 2.9 | 0.0-20.0 | -0.96 | 0.339 | 0 | 1 | 0.00 | 1.000 |
| OT11 | 0.0 | 0.0 | 0.0-0.0 | 0.4 | 2.9 | 0.0-20.0 | -0.96 | 0.339 | 0 | 1 | 0.00 | 1.000 |
| **OT12** | **15.5** | **21.1** | **0.0-80.0** | **80.4** | **90.9** | **0.0-600.0** | **-6.25** | **0.000** | **20** | **43** | **22.46** | **0.000** |
| **OT21** | **5.9** | **13.4** | **0.0-60.0** | **27.4** | **37.1** | **0.0-140.0** | **-3.68** | **0.000** | **9** | **26** | **10.84** | **0.001** |
| **OT22** | **1.4** | **5.1** | **0.0-20.0** | **7.0** | **12.8** | **0.0-60.0** | **-2.67** | **0.008** | **3** | **13** | **5.68** | **0.017** |

46 TreM types were found in canopy gaps and 44 in reference sites (Table 3). Five TreM types (small (CV11) and large woodpecker cavities (CV13), small mould trunk cavities (CV23) and small (DE11) and large sun-exposed dead branches (DE12)) were unique to canopy gaps, while three (broken branches (IN23), sap runs (OT11) and small vertebrate nests (NE12)) were unique to reference sites (Table 3). The TreM types with the highest mean density in canopy gaps were epiphytic foliose and fruticose lichens (EP32), bark pockets (BA12), bark shelters (BA11), insect galleries with small bore holes (CV51) and perennial polypores (EP12). Except for the epiphytic lichens, the densities of all these TreM types were higher in canopy gaps than in reference sites (Table 3). Canopy gaps also had higher densities of trees bearing medium-sized woodpecker breeding cavities (CV12), splintered stems (IN24), small injuries exposing cambium and sapwood (IN31) and fronds of epiphytic ferns (EP34) compared to reference sites (Table 3). The TreM types with the highest mean densities in reference sites were epiphytic foliose and fruticose lichens (EP32), small (GR11) and medium-sized cavities formed by tree roots (GR12), epiphytic bryophytes (EP31) and bark pockets (BA12). The density of trees bearing small root cavities (GR11), small (IN13) and large bark losses (IN14), small dead branches not exposed to the sun (DE13), dead tree tops (DE15), bark structure (BA21), decayed canker (GR32), fresh flows of resin (OT12), and crown (OT21) or bark microsoil (OT22) was higher in reference sites than in canopy gaps (Table 3).

### 3.4. Drivers of TreM assemblage turnover

TreM richness in canopy gaps was positively correlated with both the density of living trees and elevation, in contrast to reference sites, where it was positively correlated solely with elevation (Table 2b). TreM density increased with the density of snags in both canopy gaps and reference sites (Table 2c). TreM diversity in canopy gaps decreased with the density of living trees and elevation, whereas TreM diversity in reference sites increased with the density of living trees and decreased with the number of tree species (Table 2d).

RDA explained 28.8% of the total variance (constrained inertia = 0.09, unconstrained inertia = 0.19, p < 0.001). RDA axis 1 explained 6.1% of the total variance (p = 0.001) and showed that the abundances of bark pockets (BA12), bark shelters (BA11), perennial polypores (EP12), fronds of epiphytic ferns (EP34), small injuries exposing cambium and sapwood (IN31) and insect galleries with small bore holes (CV51) were positively related to the density of snags (p = 0.001), while the abundances of fresh resin flows (OT12), epiphytic foliose and fruticose lichens (EP32), small (GR11) and medium-sized cavities formed by tree roots (GR12), bark microsoil (OT22), bark structure (BA21) and small bark losses (IN13) were positively related to the density of living trees (p = 0.001) (Fig. 3a). RDA axis 2 explained 1.8% of the total variance (p = 0.001) and suggested that the abundances of large cleavages formed by tree roots (GR13) and epiphytic bryophytes (EP31) were positively related to the elevation (p = 0.001; Fig. 3a). Despite the differences in TreM composition between canopy gaps and reference sites, the total number of TreM types calculated for all the study plots pooled remained constant (Fig. 3b).

**Fig. 3.**
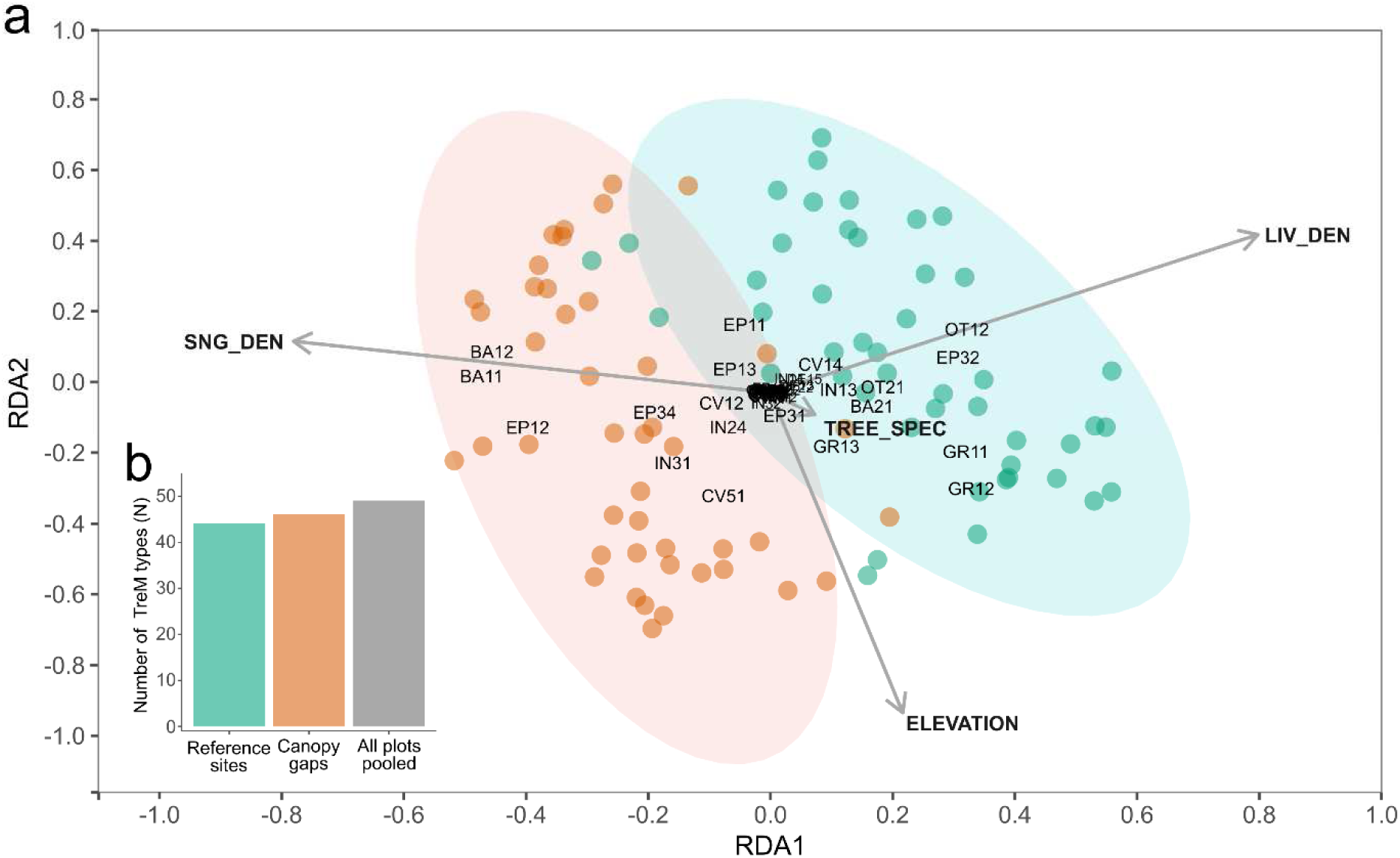
(**a**) TreM type densities (TreM codes labelled after Kraus et al. 2016; see Table S2 for the full names and descriptions of the TreM types) constrained by the effect of habitat characteristics (grey arrows) – density of living trees (LIV_DEN), density of dead standing trees (SNG_DEN), elevation (ELEVATION) and tree species richness (TREE_SPEC) – recorded in canopy gaps (copper circles) and reference sites (turquoise circles) located in Norway spruce *Picea abies*-dominated old-growth forests in the Tatra Mountains (Poland); **(b)** the total number of TreM types recorded in the reference sites (N = 46), the canopy gaps (N = 44) and all the study plots pooled (N = 90).

## 4. Discussion

Our study provides insight into the quantity and composition of TreMs in Norway spruce-dominated montane forests for patches recently disturbed by bark beetle activity and undisturbed, closed-canopy patches. Although Norway spruce is widespread and often mixed with other tree species in temperate forests, the TreM profile in natural or long-term protected single-species spruce forests has rarely been assessed. As few natural temperate Norway spruce forests have remained intact, opportunities to study TreMs in such a habitat are limited (Sabatini et al. 2018). To date, studies of TreMs on Norway spruce trees have focused on single-species managed (Asbeck et al. 2023), multispecies managed (Großmann et al. 2018; Kõrkjas et al. 2021b) and multispecies unmanaged stands (Vuidot et al. 2011; Larrieu et al. 2014). However, Zemlerová et al. (2023) have examined the natural Norway spruce forests in the Carpathians, highlighting the legacy of historical disturbances in the TreM profile. Although coniferous trees typically support fewer and less diverse TreMs compared to deciduous trees (Paillet et al. 2019), the coniferous stands usually have high tree densities, which could imply a higher abundance of distinctive TreMs (Michel and Winter 2009; Asbeck et al. 2019). Moreover, Norway spruce snags host more TreMs than living trees of this species (Przepióra and Ciach 2023). Consequently, elevated tree mortality and reduced tree density are expected to trigger major changes in the TreM assemblage in bark beetle-induced canopy gaps.

### 4.1. The role of bark beetles as ecosystem engineers in shaping TreMs

We expected that bark beetle-induced canopy gaps, with their distinctive stand structure and dominance of snags, should exhibit different TreM richness, density and diversity compared to more homogeneous closed-canopy forest. However, we found no differences in the general TreM indices between canopy gaps and undisturbed reference sites, although the TreM density in canopy gaps was marginally higher than in reference sites. The combined presence of snags and living trees leads to the most diverse stand-level TreM composition (Spînu et al. 2022). In our study, the density of living trees or snags was a significant or near-significant predictor of TreM richness and diversity in both canopy gaps and reference sites. Whereas canopy gaps created by bark beetle activity increase the abundance of snags and damaged trees, these gains may be counterbalanced by the reduced density of living trees in disturbed patches. Moreover, snags or damaged trees are often felled over time, further reducing the number of potential TreM hosts. Such stand dynamics, where numerous living trees are replaced by fewer snags, probably gives rise to the observed stable TreM richness, density and diversity – metrics that aggregate all trees and TreM types without accounting for shifts in TreM composition.

Although we found no differences in TreM indices between canopy gaps and reference sites, our results revealed substantial differences in TreM composition. Coniferous species, including Norway spruce, produce fewer large branches, have bark and wood with lower moisture retention capabilities, and exude resin, terpenes and phenolics that slow wood decomposition and cavity formation (Handegard et al. 2021). As a result, saproxylic TreMs, particularly cavities, are often rare in spruce forests (Remm and Lõhmus 2011). However, bark beetle activity triggers tree mortality and creates canopy gaps that act as local concentrations of snags, in which intensified wood decomposition processes support the formation of saproxylic TreMs. The studied canopy gaps were characterized by a higher frequency and density of TreMs such as cavities, cracks, loose bark patches and fruiting bodies of saproxylic fungi compared to a closed-canopy forest, making the bark beetle infestation patches hosts of abundant saproxylic TreMs in a single-species coniferous forest.

### 4.2. Bark beetle-driven changes in tree traits, stand structure and TreMs

In our study, the curves describing the relationship between diameter and TreM richness on living trees or snags differed between trees in canopy gaps and reference sites. The increase in TreM richness with diameter is consistent across various habitats and tree species (Paillet et al. 2019). As trees age, they accumulate damage, sustain initiated wood decay and are colonized by epiphytes, which processes lead to the formation of TreMs such as cavities, cracks, dendrotelmata and patches of mosses and lichens (Kõrkjas et al. 2021a). During canopy gap formation, additional TreMs may develop on smaller trees as a result of damage from falling trees or the activity of cambium-feeding insects. However, the open conditions of canopy gaps, where solar radiation is higher but the humidity lower, create a microclimate that slows wood decomposition and the development of certain epiphytes (Tanona and Czarnota 2022; Wu et al. 2022), potentially limiting moisture-dependent TreMs. As this study shows, large living trees and snags in canopy gaps may accumulate fewer TreMs over time compared to similar sized-trees in more humid, closed-canopy forests.

In the Białowieża Forest, TreM richness on Norway spruce trees doubled for approximately every 35 cm of growth in diameter, with a mean predicted TreM richness of 1.2 types per living tree at 10 cm diameter and 4.0 types per tree at 70 cm diameter (Przepióra and Ciach 2023); the TreM accumulation dynamics there were thus comparable to the results of this study. Norway spruces in mixed stands in the Białowieża Forest commonly hosted TreM types such as epiphytic cryptogams, root buttress cavities, bark pockets, insect galleries, sap and resin runs, fungal fruiting bodies and exposed sapwood (Przepióra and Ciach 2023), all of which were also abundant in the Norway spruce-dominated montane forest. However, the results of the present work indicate that the existence of a canopy gap is linked with the rate of TreM accumulation. These findings suggest that the TreM profile is closely associated with the tree species, but that TreM accumulation could be modified by bark beetle activity.

TreM richness in the reference sites exhibited a near-significant positive correlation with the number of tree species, whereas the correlation with TreM diversity was negative. Gaps created by local disruptions in the canopy closure allow sunlight to penetrate the stand interior, facilitating the entry of new trees to the species pool. These changes promote the regrowth of floor vegetation and tree regeneration, thereby enabling the inclusion of sun-demanding TreM-rich tree species like rowan and European aspen *Populus tremula* to the species pool (Nováková and Edwards-Jonášová 2015, Holeksa et al. 2017) and increasing the probability of the development of TreMs new to the assemblage. Regenerated post-disturbance patches represent the only sites where new tree species can reach such an age and size that novel TreMs arise (Spînu et al. 2023). On the other hand, these patches may lack large, old Norway spruces, often downed during the gap formation process (Sproull et al. 2015), potentially reducing TreM diversity. As the canopy gaps studied were relatively recent and hosted only young trees, the relationship between tree species number and TreM richness or diversity was evident only in the reference sites, where trees were able to mature. This result suggests that the relationship between tree species number and TreM richness or diversity within natural Norway spruce-dominated forests is apparent in habitat patches long after the disturbance has allowed new tree species to enter the species pool and the stand has reverted to closed-canopy forest.

### 4.3. Bark beetle disturbance as a mediator of topographic conditions influencing TreMs

We found that TreM richness increased in both canopy gaps and reference sites, while TreM diversity in canopy gaps decreased with increasing elevation. High elevations are associated with increased rainfall and humidity, which promote cavity formation and epiphyte growth (Remm and Lõhmus 2011; Asbeck et al. 2019; Kozák et al. 2023). Moreover, steep slopes are frequently encountered at high elevations, which can lead to rockfalls causing the trunk damage that supports TreM formation (Larrieu et al. 2022). However, the density of trees that potentially host TreMs drops with increasing elevation (Erschbamer and Wallnöfer 2007). Consequently, higher elevations may foster the introduction of new structures to the assemblage, thereby increasing TreM richness but decreasing tree density at higher elevations, amplified in canopy gaps as a result of tree mortality, reduce TreM diversity in upper montane forest zones.

Our study showed that TreM richness on individual trees was lower on north-facing slopes but higher on east-facing slopes in canopy gaps. In the Tatra Mts., the very strong Föhn-type winds blowing from the south, make trees on south-facing slopes more prone to damage (Ochtyra 2020), contributing to the formation of TreMs such as branch and trunk breakage. Wind exposure also promotes the development of reaction wood, which forms root buttress cavities (Crook et al. 1997; Homma 1997) and induces asymmetric crowns that are vulnerable to snowfall-induced damage (MacFarlane and Kane 2017), common in montane forests (Hess 1996). Additionally, the higher solar radiation on south– and east-facing slopes reduces humidity, which negatively affects moisture-dependent Norway spruces. Reduced canopy closure exposes individual trees to winds and solar radiation to a greater extent than trees in closed-canopy forests (Siegert et al. 2024). This stress can undermine tree health, promoting fungal growth, resin flow and wood decay, which, in turn, increase the likelihood of TreMs occurring on individual standing trees. Our results suggest that bark beetle-induced canopy gaps, weakly protected against wind and solar radiation, amplify the effects of slope exposure on TreMs found on individual trees compared to the less extreme abiotic conditions in closed-canopy forests.

The TreM richness found on individual trees decreases with terrain ruggedness, as indicated by TPI, in both canopy gaps and reference sites. Although trees on ridges or mountain summits are more susceptible to wind damage, individuals growing on slopes are vulnerable to falling rock debris and avalanches, which can damage trunks and promote the formation of TreMs such as exposed sapwood, loose bark and broken crowns (Stokes et al. 2005). Moreover, the humid microclimate in gully or valley bottoms may also favour moisture-dependent TreMs (Pouska et al. 2016; Man et al. 2022). Our results suggest that slopes or valleys provide better conditions for TreM formation than ridges or hilltops, and such favourable conditions persist even after canopy gaps have opened up.

### 4.4. Bark beetle-related TreMs and biodiversity boost

The engineering activities of bark beetles, which result in local concentrations of saproxylic TreMs, potentially support biodiversity. Previous studies in the Tatra Mts. have shown that bark beetle-induced canopy gaps have a higher richness, diversity and abundance of breeding birds, particularly insectivorous ground– and secondary cavity-nesters (Przepióra et al. 2020). Secondary cavity-nesters rely on diverse cavities, the high density of which in gaps, including unique types not occurring in a closed-canopy forest, provides a range of niches for these groups of birds (Remm et al. 2006). Moreover, the high density of trees with cracks and loose bark, splintered stems and epiphytic ferns, found in the studied gaps, provide nesting sites for species such as Eurasian treecreeper *Certhia familiaris* and Eurasian wren *Troglodytes troglodytes*, both of which are more frequent and abundant in canopy gaps than in closed-canopy forest (Przepióra et al. 2020).

TreMs formed by bark beetle activity may also be important for bats and invertebrates. The activity of the western barbastelle *Barbastella barbastellus* increases in bark beetle-infested areas (Rachwald et al. 2022), and the greater densities of bark pockets, bark shelters, cracks, splintered stems and cavities within such areas provide roosting sites, allowing the species to inhabit roost-poor coniferous forests. Moreover, tree mortality triggered by bark beetles leads to the emergence of deadwood in which saproxylic insects develop in large numbers (Müller et al. 2008). Then, the diversity of the invertebrates is further supported by TreMs such as fruiting bodies of polypore fungi and sun-exposed dead branches that offer suitable substrates for fungivorous or thermophilous species (Larrieu et al. 2018). Invertebrates developing within canopy gaps produce galleries and boreholes that serve as shelters for other organisms, thus enhancing biodiversity (Larrieu et al. 2018; Gottfried et al. 2019). Our results indicate that bark beetles, being ecosystem engineers that trigger cascading processes in the ecosystem, locally expand the TreM composition to include rare, primarily saproxylic, TreMs, essential for maintaining biodiversity in coniferous forests.

## 5. Conclusion

Our study provides a reference for the quantity and composition of TreMs in natural, Norway spruce-dominated montane forests. By comparing patches affected by bark beetles with undisturbed closed-canopy sites, we demonstrate the cascading effect of bark beetle activity on the TreM assemblage and describe the role of an ecosystem engineer in shaping TreMs in a forest ecosystem. Although the general TreM indices did not differ between the canopy gaps and undisturbed forest patches, bark beetle-induced canopy gaps led to a local turnover of the TreM assemblage, facilitating the concentration of saproxylic TreMs, which are crucial for ecosystem biodiversity but rare in closed-canopy forests. Furthermore, the results highlight differences in TreM accumulation between living and standing dead trees in gaps and undisturbed forests, potentially resulting from the local modification of topographical and microclimatic influences on TreM development caused by canopy gap formation. Our study contributes to the understanding of biodiversity dynamics in Norway spruce-dominated montane forests, highlighting the interplay between bark beetle activity, habitat characteristics and TreM numbers. Natural processes associated with insect outbreaks in coniferous forests, such as higher small-scale tree mortality, modify the TreM assemblage, and increase habitat heterogeneity and forest complexity at the landscape scale by the creation of a mosaic of disturbed and undisturbed habitat patches. These findings underscore the role of ecosystem engineers in shaping the resources essential for biodiversity in temperate forests. Our results stress the need to integrate natural disturbance dynamics into forest management strategies in order to support biodiversity in Norway spruce-dominated forests.

## Acknowledgements

This study was financially supported by The National Science Centre in Poland from the Preludium grant (2021/41/N/NZ9/03441) and the Opus grant (2021/41/B/NZ8/03456), and also by the Ministry of Science and Higher Education of the Republic of Poland within the framework of statutory funds awarded to the Faculty of Forestry, University of Agriculture. Fabian Przepióra was supported by the Foundation for Polish Science (FNP). We thank Józef Bobak and Antoni Zięba from the Tatra National Park for their assistance during the fieldwork. We are also grateful to the Institute of Geography and Spatial Organization of Polish Academy of Sciences for providing accommodation in the study area.

## Ethical statement

The study was performed in accordance with Polish law.

## Data availability

The data used in this study is available on request from the corresponding author.

## Supplementary Materials

**Table S1.**
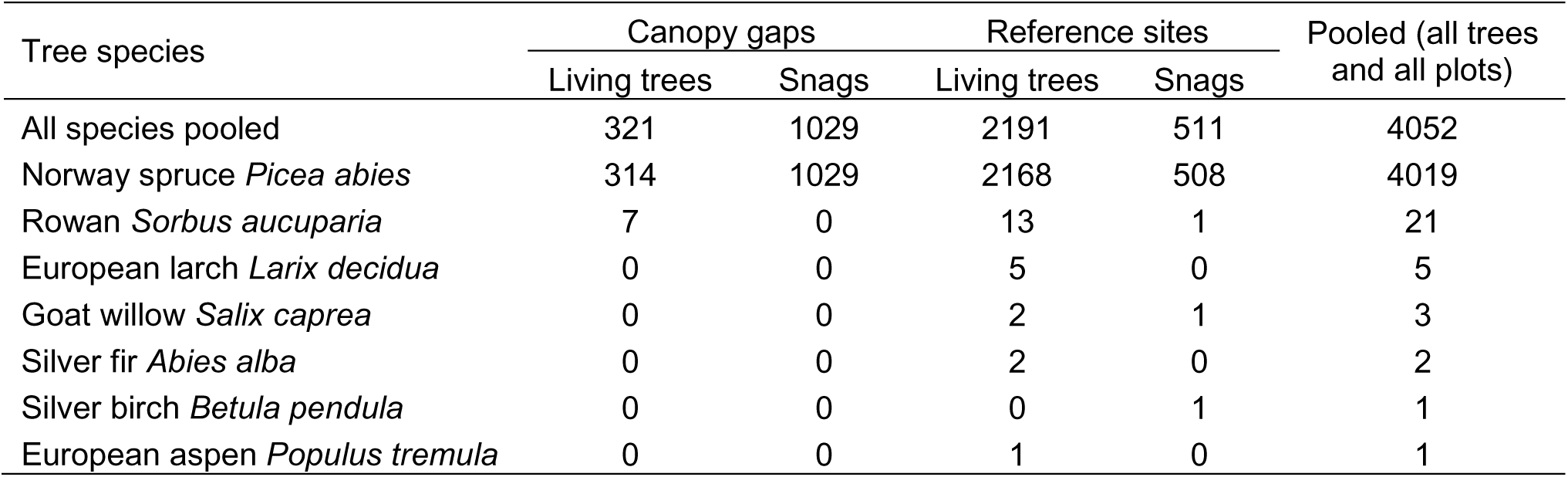
Numbers of sampled living trees and dead standing trees (snags) of Norway spruce *Picea abies*, rowan *Sorbus aucuparia*, European larch *Larix decidua*, goat willow *Salix caprea*, silver fir *Abies alba*, silver birch *Betula pendula*, European aspen *Populus tremula* and all species pooled found in canopy gaps (N = 44) and reference sites (N = 46), as well as all species and all study plots pooled in the Norway spruce-dominated old-growth montane forest in the Tatra Mountains (Poland).

**Table S2.**
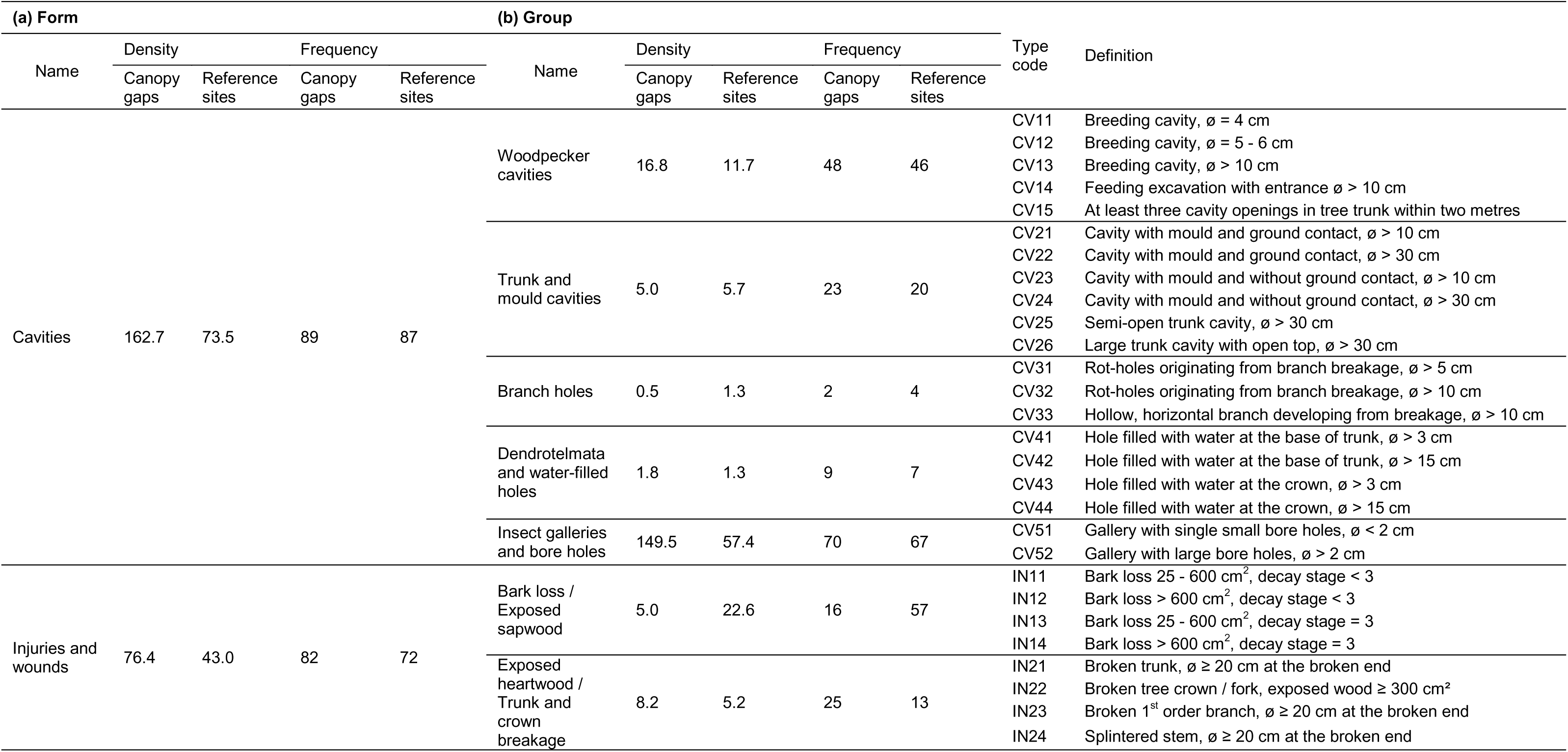
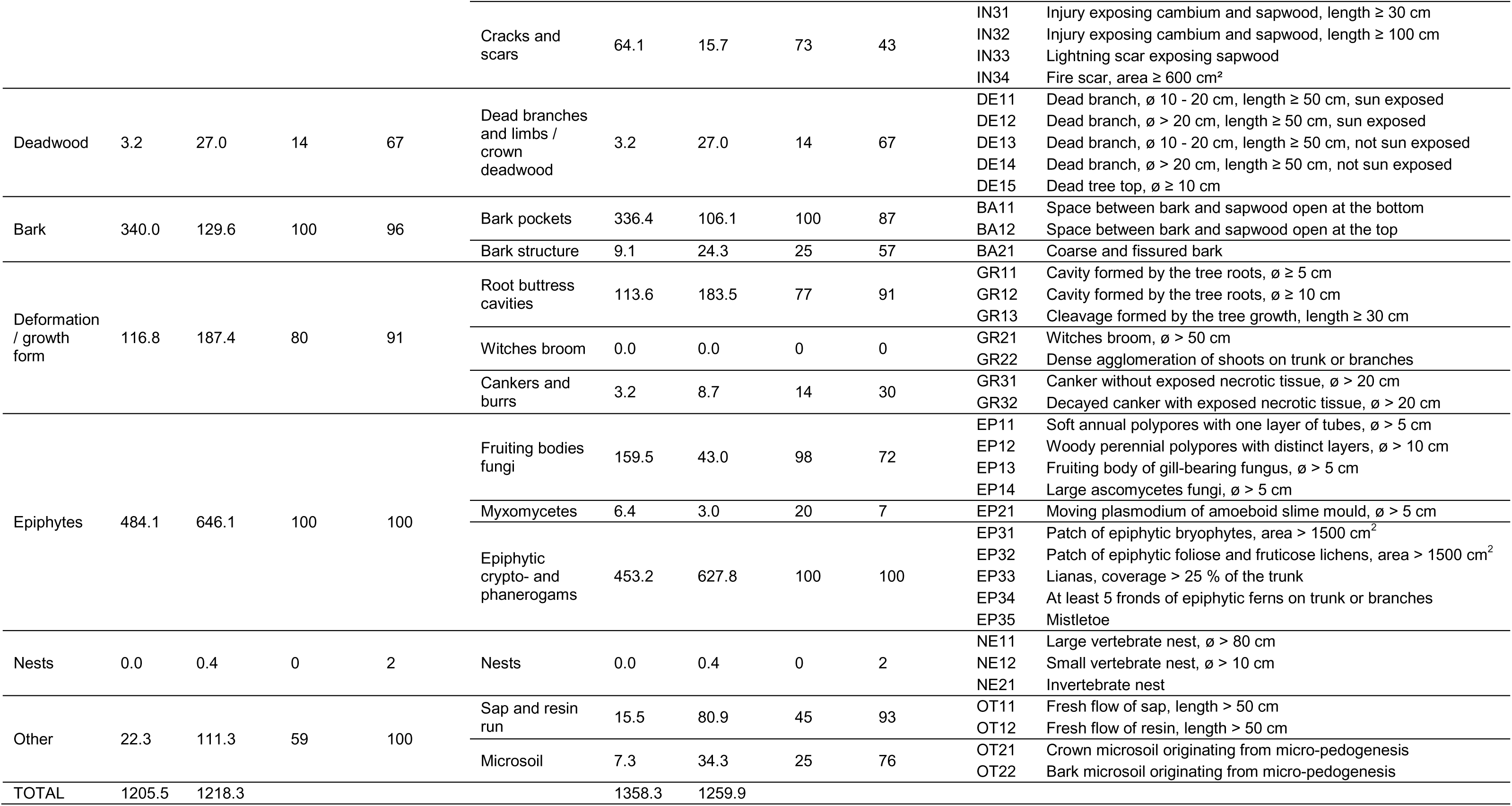
Tree-related Microhabitat (TreM) densities (density of trees bearing a given TreM ha^-1^) and frequencies (percentage of plots with a given TreM) of. **(a)** forms and **(b)** groups of TreMs found in study plots located in canopy gaps (N = 44) and reference sites (N = 46) in the Norway spruce *Picea abies*-dominated old-growth forests in the Tatra Mountains (Poland). The names, groups and codes of types of the TreM forms have been adapted from Kraus et al. (2016).

**Fig. S1.**
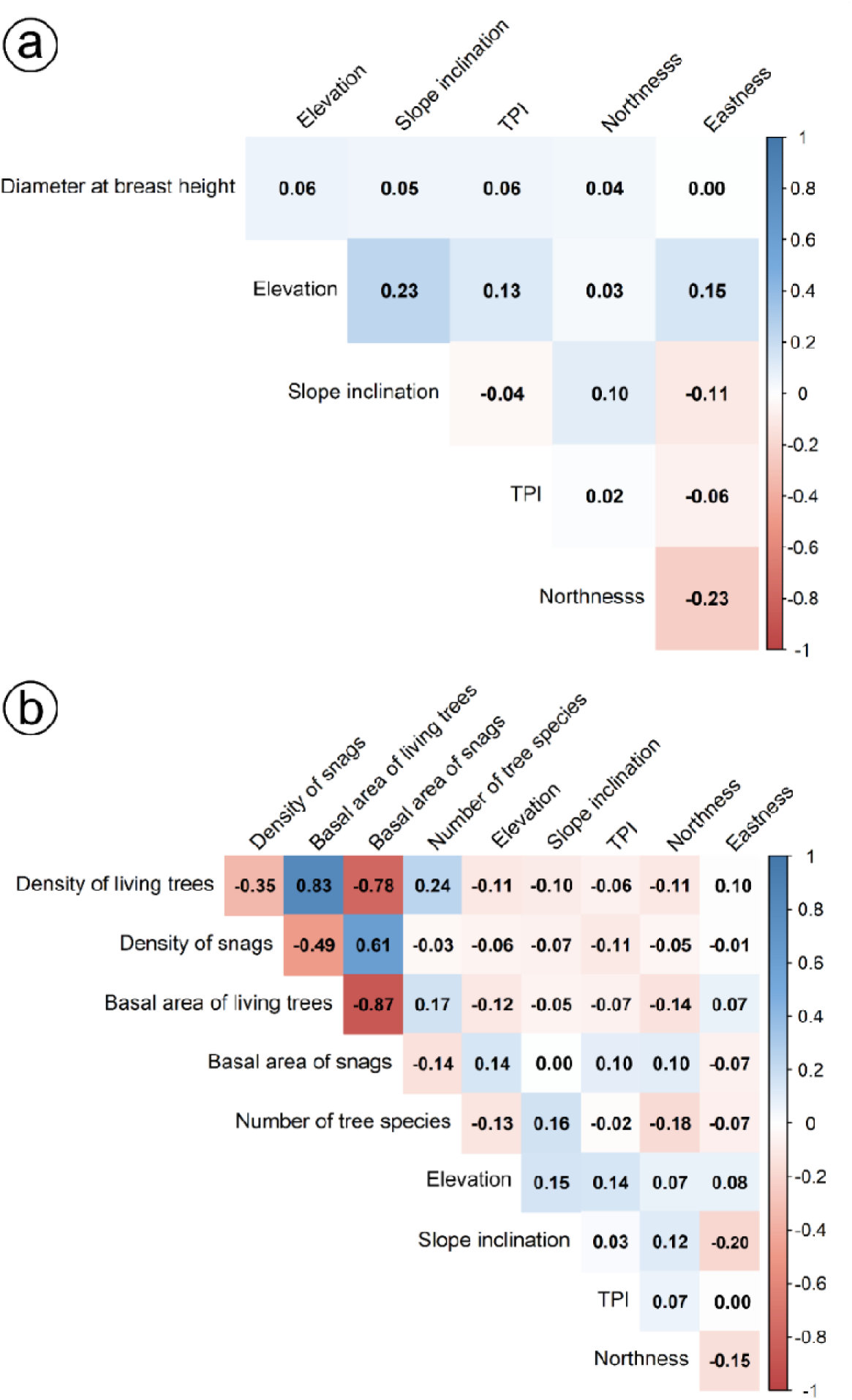
Pearson correlation coefficients between variables **(a)** at the individual tree level: diameter at breast height, elevation, slope inclination, Topographic Position Index (TPI), northness and eastness; and **(b)** at the study plot level: density of living trees, density of snags (dead standing trees), basal area of living trees, basal area of snags, number of tree species, elevation, slope inclination, TPI, northness and eastness, found on plots in the Norway spruce-*Picea abies* dominated old-growth montane forest in the Tatra Mountains (Poland).

